# SGID: a comprehensive and interactive database of the silkworm

**DOI:** 10.1101/739961

**Authors:** Zhenglin Zhu, Zhufen Guan, Gexin Liu, Yawang Wang, Ze Zhang

## Abstract

Although the domestic silkworm (*Bombyx mori*) is an important model and economic animal, there is a lack of comprehensive database for this organism. Here, we developed the silkworm genome informatics database, SGID. It aims to bring together all silkworm related biological data and provide an interactive platform for gene inquiry and analysis. The function annotation in SGID is thorough and covers 98% of the silkworm genes. The annotation details include function description, gene ontology, KEGG, pathway, subcellular location, transmembrane topology, protein secondary/tertiary structure, homologous group and transcription factor. SGID provides genome scale visualization of population genetics test results based on high depth resequencing data of 158 silkworm samples. It also provides interactive analysis tools of transcriptomic and epigenomic data from 79 NCBI BioProjects. SGID is freely available at http://sgid.popgenetics.net. This database will be extremely useful to silkworm research in the future.

## Introduction

The silkworm, *Bombyx mori*, domesticated from its wild ancestry, *B. mandarina*, nearly 5000 years ago, contributes to silk industry, pest control (Gu et al., 2017; Li et al., 2015) and evolutionary biology. It is a promising model organism in life sciences (Meng et al., 2017). Because of its importance, the silkworm genome was sequenced and annotated in 2004 (Mita et al., 2004; Xia et al., 2004) and supplementarily annotated in 2012 (Shao et al., 2012). According to the records in NCBI PubMed, more than nine thousand silkworm related works have been published. The chromosome-level assembly of the silkworm genome is accomplished in 2017 and published in 2019 (Kawamoto et al.). Meanwhile, with the development of the sequencing technology, massive silkworm transcriptomic or epigenomic data were produced (Gu et al., 2019; Li et al., 2019a; Li et al., 2019b; Wu et al., 2019; Xiang et al., 2018). Up to now, there are already more than one thousand records of silkworm DNA-seq or RNA-seq data documented in NCBI SRA database. A comprehensive archiving and synthesis of these data is important to silkworm research.

Many model organisms have their own online bioinformatics analysis platform, such as TAIR for *Arabidopsis* (Poole, 2007), Flybase (Ashburner and Drysdale, 1994; Thurmond et al., 2018) for *Drosophila* and MGI (Bult et al., 2019; Smith et al., 2019) for mouse. Currently, SilkBase (Mita et al., 2003) is the only workable dedicated database for silkworm. It mainly focuses on the archive of sequences and is lack of analysis tools. Ensembl Silkworm (Kersey et al., 2018; Zerbino et al., 2018) and InsectBase (Yin et al., 2016) are absence of update and without workable whole genome browser. Until now, there is still not a comprehensive online analysis platform of the silkworm.

To assist silkworm research, we developed the silkworm genome informatics database, SGID, through collecting and cataloguing comprehensive genomics, transcriptomics, proteomics and epigenomics data. On the basis of previous works (Duan et al., 2010; Kawamoto et al., 2019; Mita et al., 2003; Wang et al., 2005), we thoroughly annotated silkworm genes in the contents of function, protein structure, homolog and transcription factor. We also incorporated repeat elements, population statistic tests and epigenomic analysis results into the genome browser, which will help users to get a more comprehensive picture of genome segments. We developed interactive and click-one type analysis tools in SGID, letting users to obtain one or more genes’ overall information swiftly.

## Materials and methods

### Data processing for basic gene annotation

We used the high quality assembly of the silkworm genome (Kawamoto et al., 2019) as the reference. Based on gene models (2017) in SilkBase, we re-annotated all silkworm genes by UniProt (Apweiler et al., 2004; Patient et al., 2008). Generally speaking, we BLAST the protein sequence of each gene against the UniProt protein database and took hits with significant similarity (*E-value* < 0.05, Coverage > 0.7). In this way, we made connections of the silkworm genes and UniProt proteins, and obtained UniProt annotations of the silkworm genes, including Pubmed ID, EMBL ID, Proteomes ID, Pfam (Bateman et al., 2000; Finn et al., 2016), Interpro (Mitchell et al., 2019; Mulder et al., 2002), Gene Ontologies (GO), KEGG (Kanehisa, 2002; Kanehisa et al., 2017) and so on.

To validate gene expression in protein level, we manually collected all sequenced silkworm peptides referred in published proteomics related works and did alignments of them and all silkworm genes. We filtered out results with cutoffs of *E-value* <0.05 and coverage > 0.8. If the predicted protein of one gene matches two or more peptides, we consider this gene has expression evidence.

We used CD-HIT (Li and Godzik, 2006) to search for homologous genes, requiring identify > 50% and coverage > 70%, and identified 1064 gene clusters. We did multiple alignments of the sequences within each cluster by MUSCLE (Edgar, 2004a; Edgar, 2004b), and built the phylogenetic tree by FastTree 2.1 (Price et al., 2010), with the parameter ‘-boot 5000’ to test trees’ likelihoods. We called repeat elements by RepeatMasker (Tarailo-Graovac and Chen, 2009; Tempel, 2012) and predicted transcription factors by the pipeline referred in AnimalTFDB 3.0 (Hu et al., 2015; Zhang et al., 2012; Zhang et al., 2019).

### Subcellular localization and structure prediction

Annotations from UniProt only cover part of the silkworm proteins. Thus, we re-do protein function and structure prediction for all genes. We predicted subcellular locations of ASFV proteins through CELLO v2.5 (Yu et al., 2006; Yu et al., 2004) and transmembrane helixes within proteins’ sequences using TMHMM 2.0 (Krogh et al., 2001). The output images were converted into PNG format for display in websites by Magick (www.imagemagick.org).

We BLAST all protein sequences against the PDB database (Berman et al., 2000; Burley et al., 2019) and extracted significant alignment results as we did for UniProt. We also did protein structure prediction by InterproScan (Jones et al., 2014; Zdobnov and Apweiler, 2001) and put significant results into SGID. We used SignalP (Almagro Armenteros et al., 2019; Nielsen, 2017) to predict signal peptides for silkworm proteins.

### Gene Ontolgy

We put the GO (Gene Ontology) from InterproScan, UniProt and SilkBase together as SGID GO data set. We made alignments of the silkworm genes and KEGG silkworm proteins, and extracted the hits with *E-value* <0.05 and identity >0.9. In this way, we made connections between the silkworm genes and KEGG proteins. We also extracted KEGG pathway information (Aoki-Kinoshita and Kanehisa, 2007) from KEGG and made connections between pathways and silkworm genes using KEGG protein ID as a bridge. We obtain Entrez IDs of silkworm genes by KOBAS (Wu et al., 2006; Xie et al., 2011).

To enable a gene search using old gene models (Wang et al., 2005; Xia et al., 2004), we made connections between old gene models and SilkBase gene models 2017. Like we did for KEGG proteins, we made alignments of predicted proteins between old gene models and gene models 2017 and selected the best hit of each alignment.

### Pre-processing of transcriptomic and epigenomic data

We collected transcriptomic and epigenomic data of silkworm related projects from NCBI. For transcriptomes, we classified them into three categories, ‘DEG’ (differentially expressed genes), ‘Stage’ and ‘Tissue’. ‘DEG’ means the project is to identify differentially expressed genes in different experimental conditions. ‘Stage’ means the project is to observe gene expression at stages of different time points. Tissue means the project is to obtain expression profiling in different tissues. Following the standard RNA-seq analysis protocol (Ghosh and Chan, 2016; Pollier et al., 2013), we mapped transcriptomic reads onto the reference genome by bowtie (Langdon, 2015; Langmead and Salzberg, 2012) and called FKPM (Fragments per Kilobase Million) by cuffnorm (Wang et al., 2017).

For epigenomic data, according to experimental methodologies, we classified them into ChIP-Seq, Bisulfite-Seq and miRNA. ChIP-Seq stands for combining chromatin immunoprecipitation (ChIP) assays with next-generation sequencing. Bisulfite-Seq is the use of bisulfite treatment of DNA before routine sequencing to determine the pattern of methylation (Chatterjee et al., 2012). Small RNA means small RNA sequencing. For ChIP-Seq data, we used Bowtie to align reads onto the reference genome and inspected signatures by MAC2 (Liu, 2014; Zhang et al., 2008). We used Bismark (Krueger and Andrews, 2011) to pre-process Bisulfite-Seq data. We aligned small RNA reads onto the reference genome by Bowtie. Epigenomic analysis results are converted into bigWig format by the UCSC Genome tool bedGraphToBigWig for display in terminals.

### Identification of domestication genes

We used Bowtie to map the genome resequencing data of 142 domesticated and 16 wild silkworm samples, including PRJDB4743, PRJNA402108 and an unpublished resequencing data (depth = 30x) of 15 silkworm samples produced by our lab, onto the reference genome and made bam files by SAMtools (Li et al., 2009). To illustrate the evolution pressure at a genome wide scale, we slid along the silkworm genome with a window size of 2000 bp and a step size of 200 bp. In each sliding window, we calculated Pi, Theta (Watterson, 1975), Tajima’s D (Tajima, 1989) and the composite likelihood ratio (CLR) (Nielsen et al., 2005) by ANGSD (Durvasula et al., 2016; Korneliussen et al., 2014) and SweapFineder2 (DeGiorgio et al., 2016). We also did the four population genetics test for each gene and made coalescent simulation (Pavlidis et al., 2010) ranking test (CSRT) (Zhu et al., 2007) according the silkworm domestication mode (Yang et al., 2014). We called domestication genes in a strict method, requiring a Tajima’s D _domesticated_ < −1, a CSRT <0.05, a Tajima’s D _domesticated_ < Tajima’s D _wild_, a Tajima’s D _min, domesticated_ > top 5% point value in ascending order, a Fst _max_ > the top 5% point value in descending order and a CLR _max, domesticated_ > the top 5% threshold in the whole genome. ‘Min’ or ‘max’ indicates the minimal or max value in the genic region extended by 10% of gene length, to accommodate the situation that the 5’ or 3’ terminals of a gene is under evolutionary forces. A subscript of ‘domesticated’ or ‘wild’ means the population genetics test is done on domesticated or wild silkworms. We also appended identified domestication genes in (Xia et al., 2009) and (Xiang et al., 2018) to our domestication genes dataset. We searched for genes possibly under balancing selection in the criteria that a Tajima’s D _domesticated_ > the top 5% point value in ascending ranking, a Tajima’s D _wild_ < 0.5, a Tajima’s D_domesticated_ >1 and a CSRT < 0.95.

### Genome browser and analysis tools

The genome browser of SGID is developed based on an open source population genetics visualization and analysis package SWAV (swav.popgenetics.org). We used MSAViewer (Yachdav et al., 2016) to show multiple alignments of homologous proteins, and phylotree (Shank et al., 2018) to display phylogenetic trees of gene clusters. The fuzzy text search in the home page is compatible with gene ID, gene name and gene function annotations. The alignment search tools in SGID are Perl codes to parse BLAT (Kent, 2002) or NCBI BLAST results (Johnson et al., 2008). The interface to exhibit gene expression is built upon D3 and JQuery. The overall web structure is Mysql + PHP + CodeIgniter (www.codeigniter.com) + JQuery (jquery.com).

## Result and discussions

### The biological data in SGID

SilkBase annotated 16880 genes in the high quality assembly of the silkworm genome (Kawamoto et al., 2019), but left 3329 without function descriptions. For these, SGID incorporated protein information from UniProt (Apweiler et al., 2004; Patient et al., 2008) and re-annotated the functions of 15594 genes, within which 2962 got function annotations for the first time. For a lot of genes, SGID gives not only simple descriptions, but also information on function details, chemical properties, related publications, protein structure, topologies, pathways and gene ontologies. In addition to the available gene ontology (GO) annotations of 9147 genes in SilkBase, SGID newly labeled GO IDs for 5521 genes. Besides, SGID made KEGG annotations for 16028 genes and Entrez IDs for 16320 genes. These are important for research, especially for gene set function enrichment analysis (Fig. 1). Using peptide sequences from published experiments, we validated 2999 protein coding genes. They are of proteomics evidence. To depict one gene’s function in a cell, SGID provides information on gene’s subcellular localization and topology prediction. More than half (9592, 56.8%) of the silkworm genes are located in the nuclear, and 2878 genes (17.0%) have transmembrane regions (Fig. 2). Furthermore, 1960 silkworm genes are predicted to have signal peptides. Encouragingly, 9844 silkworm proteins are of PDB matches with *E-value* <0.05, which infers that more than half (58.3%) silkworm expressed proteins have structural information. External links to UniProt Proteomes, PRIDE (Perez-Riverol et al., 2019; Reisinger et al., 2015), Pfam (Bateman et al., 2000; Finn et al., 2016), Interpro (Jones et al., 2014; Zdobnov and Apweiler, 2001), SUPFAM (Pandit et al., 2004), Gene 3D (Dawson et al., 2017; Pearl et al., 2003; Pearl et al., 2002), Protein Modal Potal (Arnold et al., 2009; Haas et al., 2013) and PANTHER (Mi et al., 2017; Mi and Thomas, 2009) are also provided and they are helpful to understand the protein structure and related functions of one gene.

**Figure 1.**
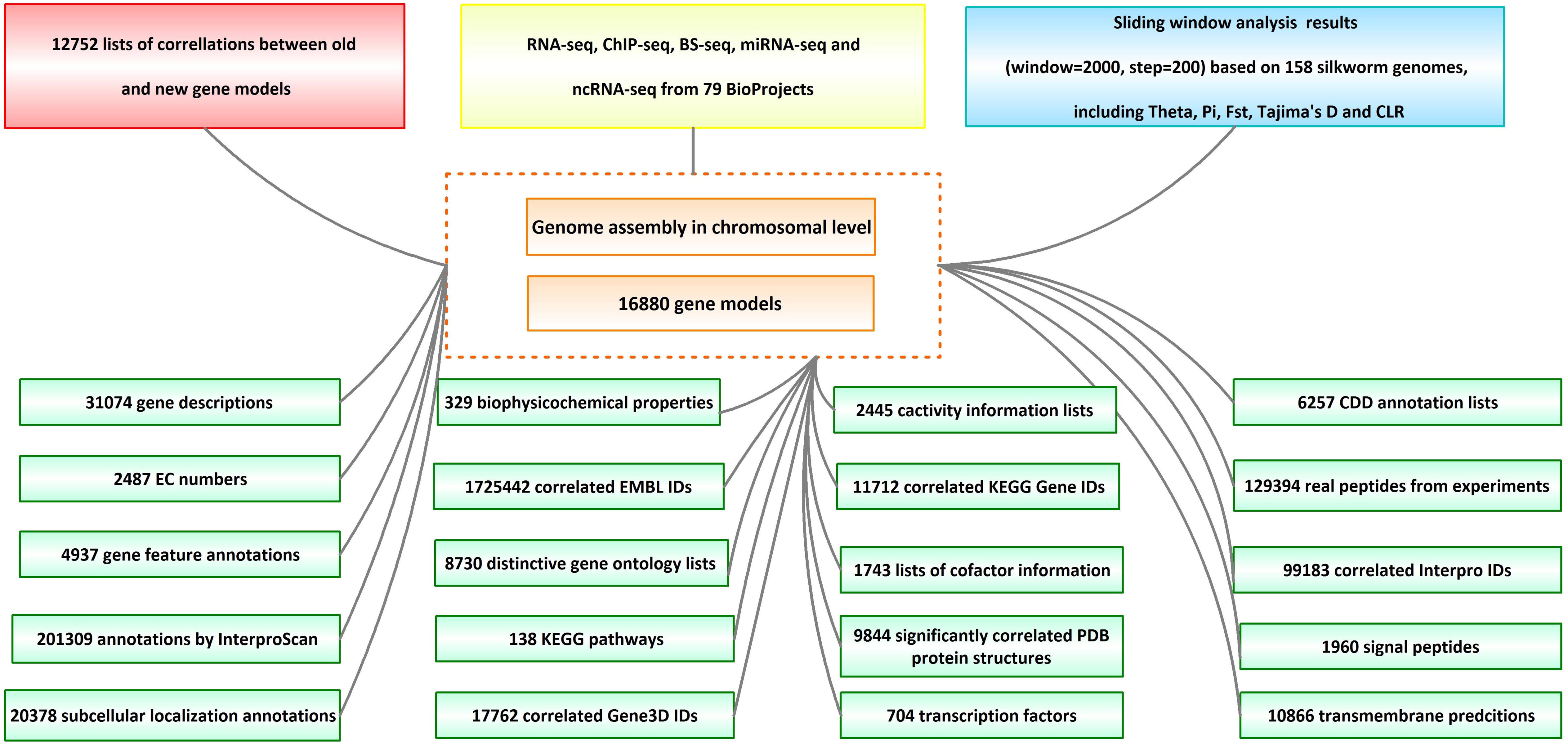
An overview of the data in SGID.

**Figure 2.**
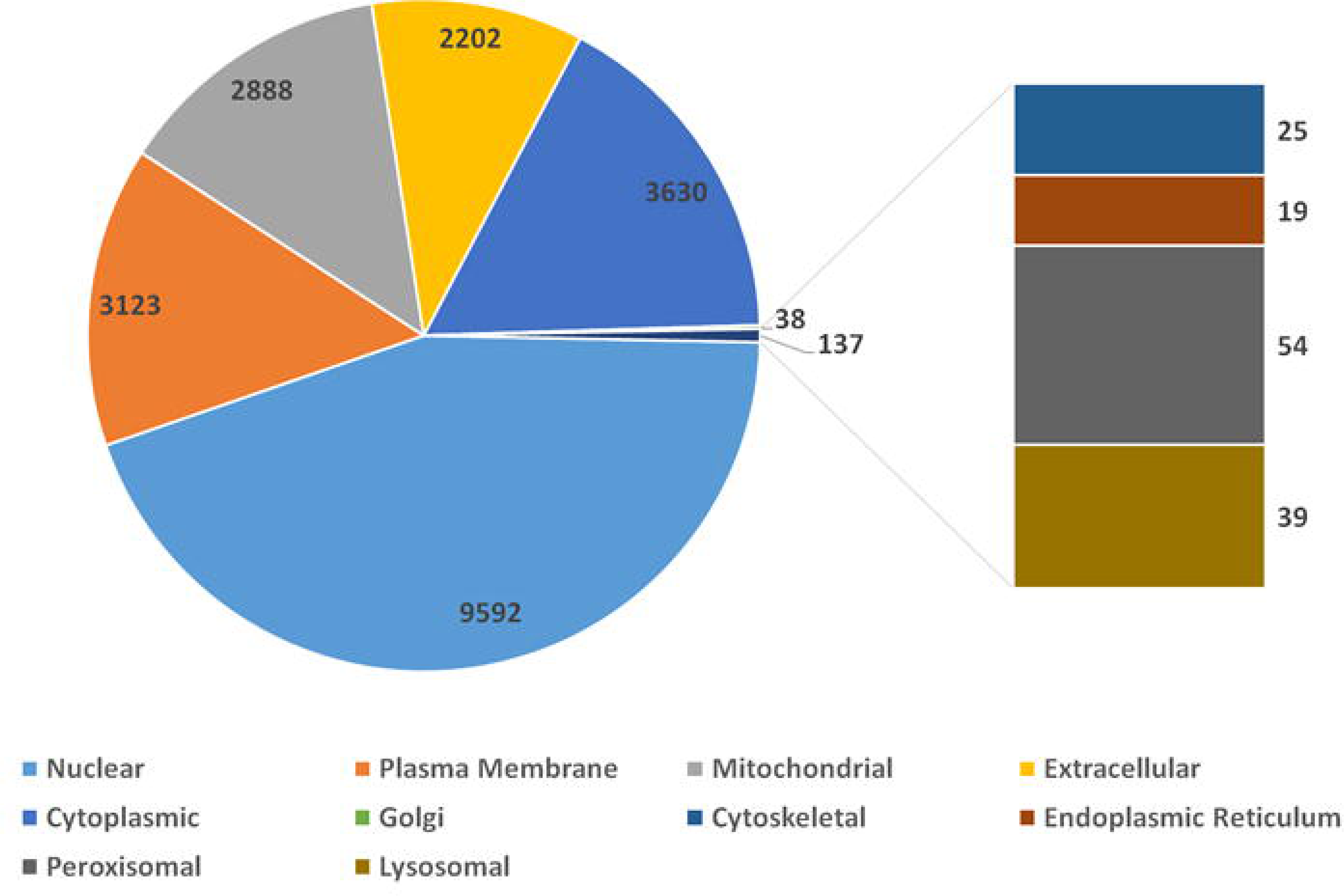
The distribution of silkworm genes in different cellular subunits.

As a domesticated insect, the silkworm is important in evolution research. Totally, we identified 569 domestication gene candidates. Users can view and inspect theses domestication genes by a SGID tool named “Population Genetics”. Population genetics test results are also displayed in the genome bowser, where users can do sliding window analysis of interested genomic or genic segments. We also identified 81 genes possibly under balancing selection.

SGID includes transcriptomic data of 41 projects and epigenomic data of 38 projects. For transcriptomes, 28 are ‘DEG’, 9 are ‘Stage’ and 4 are ‘Tissue’ as we described in Materials and Methods. SGID includes 704 transcription factors (TF) belonging to 68 TF families. It also has 571401 repeat segments covering 27.5% of the silkworm genome, which is generally in accordance with previous records (Osanai-Futahashi et al., 2008). There are more retrotransposons (93%) than DNA transposons (7%). For retrotranposons, most are LINE (46%) and SINE (44%).

### The genome browser in SGID

In SGID’s genome browser page (Fig. 3), users can view the silkworm genes, repeat elements and population genetics test tracks subsequently. An input box and a list of buttons above the browser allow users to move, zoom in, zoom out, setting focus bar, generating figures or downloading the data of one track. A click onto a gene figure will take users to the gene detail page. Clicking on one point of some track will raise a dialog displaying the value at the point. Except for a genome browser, SGID also provides a browser to view epigenomic data. In the browser, users could view gene regulation signals at some specific genome position.

**Figure 3.**
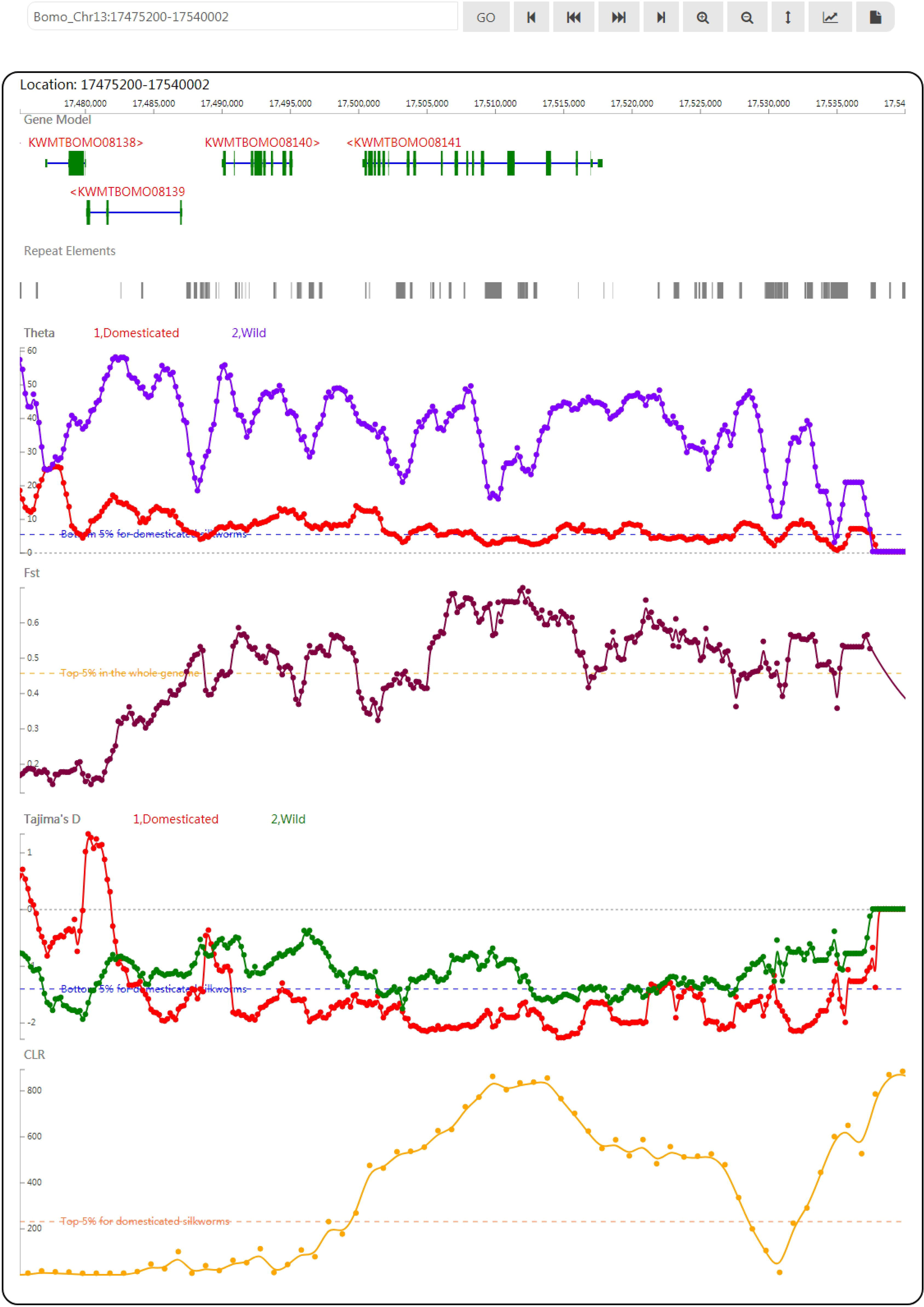
A snapshot of the genome browser of SGID with KWMTBOMO08141 in the center. Below are tracks of population genetics test results.

### Retrieve genes’ information from SGID

As a one-click type platform, SGID offers to search genes by a gene ID, a gene name, a gene function or even a brief description. In the page displaying search results, there are a list of gene information buttons within each result list. With the buttons, users can jump to to view gene details, a gene in genome browser, gene ontology and pathway, gene expression, regulation elements, gene structure and population genetics analysis results (Fig. S1A). In the detail page of each gene, aside from basic annotations (such as gene name, description, subcellular location and sequences, Fig. S1B), six information groups are listed subsequently, including “Summary”, “Ontologies”, “Topology”, “Population Genetics”, “Multiple Alignment” and “Gene Tree”. “Summary” mainly includes information resulted from protein sequence analysis (Fig. S1C). “Ontologies” displays a gene’s annotation on GO, KEGG Function, KEGG Pathway, and PANTHER (Fig. S1C). In the part of “Topology”, transmembrane regions are listed and marked in a diagram (Fig. S1F). If one gene’s protein product is of signal peptide, the region of the signal peptide will also be listed and marked. “Population Genetics” listed 5 population genetic test results (Pi, Theta, Tajima’s D, CLR and CSRT) and will give an interpretation about evolutionary forces. “Multiple Alignment” and “Gene Tree” displayed the multiple alignment of homologous genes at protein level and the phylogenetic tree produced based on the alignment.

To facilitate users to analyze a list of genes, SGID also offers to generate a list of gene information buttons through inputting a list of gene IDs. With the buttons, users can jump to some information view page directly like they do in search result page as referred above. Analogously, users can input a list of chromosome positions and obtain a list of genomic infomration links, with which users can view the genome browser or the epigenomics browser swiftly.

### SGID analysis tools

To help users to visit data more quickly, we developed a list of analysis tools in SGID. As shown in the home page, “Gene Ontology” is a tool to retrieve GO, KEGG or Entrez numbers using a list of gene IDs. “Transcriptome” is a tool to view the expression of several genes in different experiment conditions, tissue or development stages. The results will be displayed in a heatmap figure. Stopping the mouse cursor at one cell of the heatmap will display the FKPM value of one gene at an experiment condition. The project’s name is listed at the top right and users can click it to view the project’s description. “Protein Structure”, “TF”, “Population Genetics”, “Repeat Elements” and “Subcellular localization” are interactive search tools, with which users can obtain a group of genes or items with some similar biological properties. “Cluster” listed the 1064 gene clusters we identified. A summary of SGID search engines and analysis tools is shown in Fig. 4.

**Figure 4.**
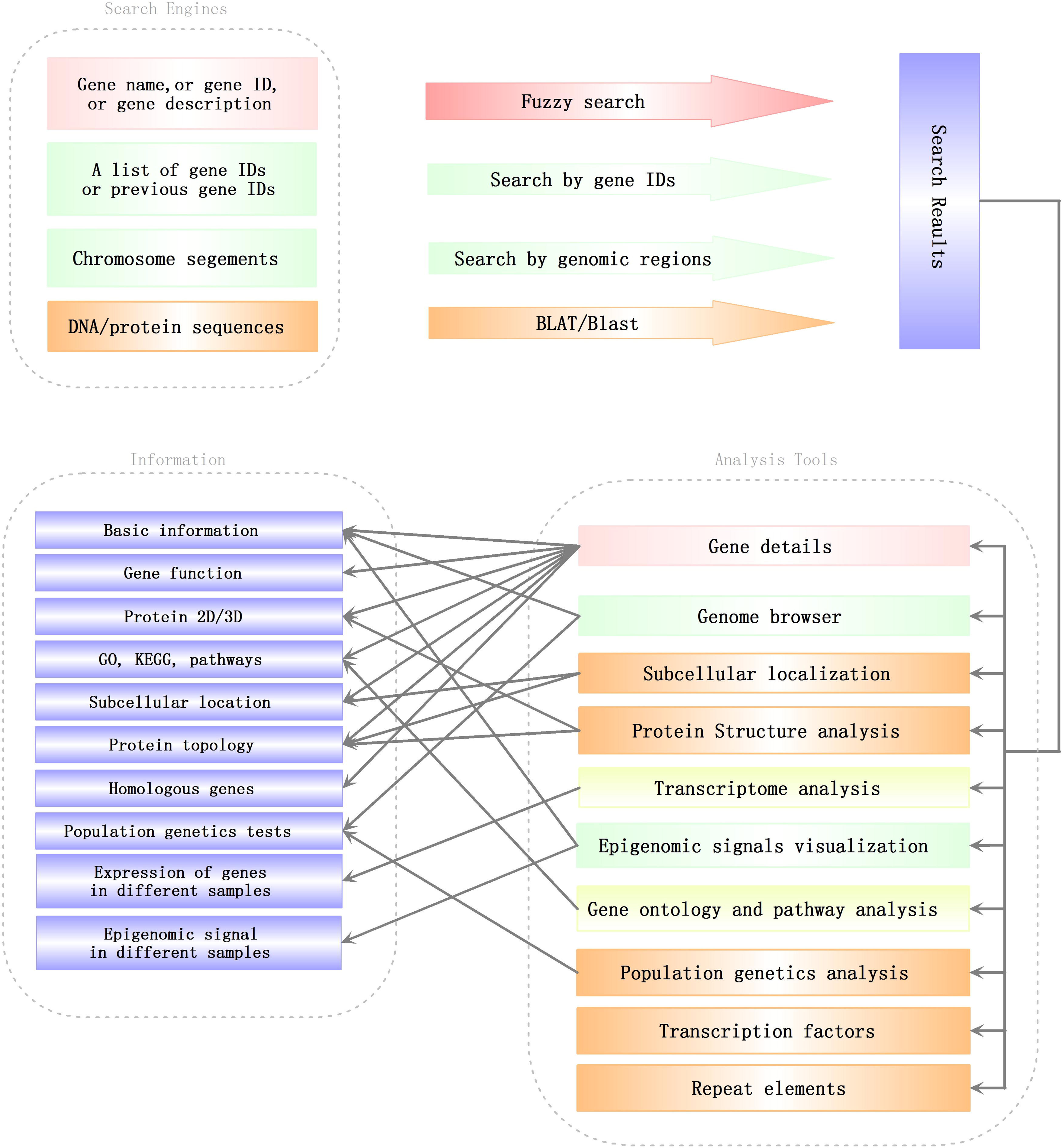
Search engines and analysis tools in SGID. Lines with arrows pointed out a general analysis flow in SGID.

### KWMTBOMO08141, a case study

KWMTBOMO08141, BGIBMGA001085 in the previous annotation (Mita et al., 2004; Xia et al., 2004), is a domestication gene referred in (8), Bmor_03834. Through searching in the home page, we found this gene is of other 17 functional related members as transient receptor (Fig. S1A). Using the buttons listed below genes in the search result page, we can obtain genes’ information one by one. In the detail page, we found this gene is of a full name “transient receptor potential-gamma protein” (Sanyal et al., 2004; Selbie et al., 1997; Woodard et al., 2007) and an alternative name “Transient receptor potential cation channel gamma”. The gene is annotated to be located in plasma membrane (Fig. S1B) and function in interacting preferentially with trpl and to a lower extent with trp (Fig. S1C). Encouragingly, the protein product of this gene is validated by experiment peptides (Fig. S1B) and of significant similarity to a real protein structure 5Z96 recorded in PDB (Fig. S1D). In gene ontology, we obtained the GO and KEGG IDs of this gene and found KWMTBOMO08141 plays roles in the pathway of phototransduction in cell membrane (Fig. S1E). In topology, this gene has 6 transmembrane regions (Fig. S1F), which is in accordance to its subcellular localization prediction. As a domestication gene, KWMTBOMO08141 has low Tajima’s D (−1.949945) and high CLR (629.851816). In the genome browser, we observed a CLR peak at the gene’s region (Fig. 2) and a higher CLR peak at the right, indicating there may be genetic hitchhiking effects in this case (Fig. S1G). In transcription analysis, we found the expression of this gene is higher in brain than other tissues (Fig. S1H) and affected by ectopic expression of ecdysone oxidase (Fig. S1I) Through scanning this gene in the SGID epigenomics browser, we observed that epigenomic signals within the genic region disappear in some cell lines (Fig. S1J).

## Conclusion

SGID is informative and user friendly. Under the idea of ‘Click-one’, SGID integrated different biological data and made them connective. SGID allows to search genes in fuzzy mode and to do analysis of more than one genes simultaneously. SGID pre-analyzed available transcriptomic data and developed a search tool to view the expression of genes in different conditions. Similar such SGID tools made the initial bioinformatics analysis of silkworm projects more efficiently. With the advance of sequencing and experiments of the silkworm, more and more data will be incorporated into SGID, making the platform to be more and more powerful.

## Data availability

All SGID data are publicly and freely accessible at http://sgid.popgenetics.net. Feedback on any aspect of the SGID database and discussions of silkworm gene annotations are welcome by email to.

## Author contributions

Z.L.Z. developed the web interface of the database. Z.Z.L., Z.G., G.L. and Y.W. collected and compiled the data and performed the analysis. Z.L.Z. and Z.Z. wrote the manuscript, conceived the idea and coordinated the project.

## Funding

This work was supported by grants from the National Natural Science Foundation of 31772524 to Z.Z., 31200941 to Z.L.Z.) and the Fundamental Research Funds for the Central Universities (106112016CDJXY290002).

## Acknowledgements

We thank Dr. Yong Zhang for insightful suggestions and Dr. Anyuan Guo for help on transcription factor prediction. We also thank Dr. Quanyou Yu, Dr. Wei Sun and Mr. Yun Wang for useful discussions.

