## Supplementary Figure S1 for "SGID: a comprehensive and interactive database of the silkworm"

A.

Bomyx mori ▾ | Transient receptor | SEARCH

Found: **18** items

| Gene | Short Name | Full Name | Annotation | Function |
|---|---|---|---|---|
| KWMTBOMO00115 (BGIBMGA002131) | dTRPA1 | Transient receptor potential cation channel subfamily A member 1 Aspartate aminotransferase | transient_receptor_potential_cation_channel_subfamily_A_member_1_[Bombyx_mori] | Essential for thermotaxis by sensing environmental temperature. Receptor-activated non-selective cation channel involved in detection of sensations such as temperature. Involved in heat nociception by being activated by warm temperature of about 24-29 degrees Celsius. |
| Gene Information · Genome Browser · Gene Ontology and Pathway · Transcriptional Analysis · View Epigenomics Data · Protein Structure · Population genetics | | | | |
| KWMTBOMO00118 (BGIBMGA002131) | dTRPA1 | Transient receptor potential cation channel subfamily A member 1 Aspartate aminotransferase | transient_receptor_potential_cation_channel_subfamily_A_member_1_[Bombyx_mori] | Essential for thermotaxis by sensing environmental temperature. Receptor-activated non-selective cation channel involved in detection of sensations such as temperature. Involved in heat nociception by being activated by warm temperature of about 24-29 degrees Celsius. |
| Gene Information · Genome Browser · Gene Ontology and Pathway · Transcriptional Analysis · View Epigenomics Data · Protein Structure · Population genetics | | | | |
| KWMTBOMO07933 (BGIBMGA001166) | | Transient receptor potential cation channel protein painless | transient_receptor_potential_cation_channel_protein_painless_[Bombyx_mori] | Receptor-activated non-selective cation channel involved in detection of pain sensation due to high temperature. Involved in heat nociception by being activated by noxious temperature of 38 degrees Celsius. |
| Gene Information · Genome Browser · Gene Ontology and Pathway · Transcriptional Analysis · View Epigenomics Data · Protein Structure · Population genetics | | | | |
| KWMTBOMO08141 (BGIBMGA001085) | TRPgamma | Transient receptor potential-gamma protein | PREDICTED:_transient_receptor_potential-gamma_protein_[Bombyx_mori] | A light-sensitive calcium channel that is required for inositide-mediated Ca(2+) entry in the retina during phospholipase C (PLC)-mediated phototransduction (By similarity). Forms a regulated cation channel when heteromultimerized with trpl. |
| Gene Information · Genome Browser · Gene Ontology and Pathway · Transcriptional Analysis · View Epigenomics Data · Protein Structure · Population genetics | | | | |
| KWMTBOMO11306 (BGIBMGA001988) | | Transient receptor potential cation channel trpm | PREDICTED:_transient_receptor_potential_cation_channel_trpm_isoform_X2_[Bombyx_mori] | Calcium channel mediating constitutive calcium ion entry. |
| Gene Information · Genome Browser · Gene Ontology and Pathway · Transcriptional Analysis · View Epigenomics Data · Protein Structure · Population genetics | | | | |
| KWMTBOMO11309 (BGIBMGA001987) | | Transient receptor potential cation channel trpm | hypothetical_protein_KGM_11096_[Danaus_plexippus] | Calcium channel mediating constitutive calcium ion entry. |
| Gene Information · Genome Browser · Gene Ontology and Pathway · Transcriptional Analysis · View Epigenomics Data · Protein Structure · Population genetics | | | | |
| KWMTBOMO15232 | | Transient receptor potential channel pyrexia | transient_receptor_potential_channel_pyrexia_[Bombyx_mori] | Receptor-activated non-selective cation channel involved in protection or tolerance from high temperature stress. Activated by temperatures above 40 degrees Celsius. More permeable to K(+) than to Na(+). May act in stress protection allow flies to survive in natural environments. |
| Gene Information · Genome Browser · Gene Ontology and Pathway · Transcriptional Analysis · View Epigenomics Data · Protein Structure · Population genetics | | | | |
| KWMTBOMO15233 (BGIBMGA005170) | | Transient receptor potential channel pyrexia | transient_receptor_potential_channel_pyrexia_[Bombyx_mori] | Receptor-activated non-selective cation channel involved in protection or tolerance from high temperature stress. Activated by temperatures above 40 degrees Celsius. More permeable to K(+) than to Na(+). May act in stress protection allow flies to survive in natural environments. |
| Gene Information · Genome Browser · Gene Ontology and Pathway · Transcriptional Analysis · View Epigenomics Data · Protein Structure · Population genetics | | | | |
| KWMTBOMO03763 (BGIBMGA010027) | | | transient_receptor_potential_channel_pyrexia_[Bombyx_mori] | |
| Gene Information · Genome Browser · Gene Ontology and Pathway · Transcriptional Analysis · View Epigenomics Data · Protein Structure · Population genetics | | | | |
| KWMTBOMO03766 (BGIBMGA010059) | | | transient_receptor_potential_channel_pyrexia-like_[Bombyx_mori] | |
| Gene Information · Genome Browser · Gene Ontology and Pathway · Transcriptional Analysis · View Epigenomics Data · Protein Structure · Population genetics | | | | |
| KWMTBOMO08346 (BGIBMGA009272) | | | PREDICTED:_transient_receptor_potential_protein_isoform_X1_[Bombyx_mori] | |
| Gene Information · Genome Browser · Gene Ontology and Pathway · Transcriptional Analysis · View Epigenomics Data · Protein Structure · Population genetics | | | | |
| KWMTBOMO08347 (BGIBMGA009272) | | | PREDICTED:_transient_receptor_potential_protein_isoform_X1_[Bombyx_mori] | |
| Gene Information · Genome Browser · Gene Ontology and Pathway · Transcriptional Analysis · View Epigenomics Data · Protein Structure · Population genetics | | | | |
| KWMTBOMO08348 (BGIBMGA009272) | | | PREDICTED:_transient_receptor_potential_protein_isoform_X1_[Bombyx_mori] | |
| Gene Information · Genome Browser · Gene Ontology and Pathway · Transcriptional Analysis · View Epigenomics Data · Protein Structure · Population genetics | | | | |
| KWMTBOMO09165 (BGIBMGA003369) | | | PREDICTED:_transient_receptor_potential_cation_channel_subfamily_V_member_5_[Papilio_polytes] | |
| Gene Information · Genome Browser · Gene Ontology and Pathway · Transcriptional Analysis · View Epigenomics Data · Protein Structure · Population genetics | | | | |
| KWMTBOMO09166 (BGIBMGA003369) | | | PREDICTED:_transient_receptor_potential_cation_channel_subfamily_V_member_5_[Papilio_xuthus] | |
| Gene Information · Genome Browser · Gene Ontology and Pathway · Transcriptional Analysis · View Epigenomics Data · Protein Structure · Population genetics | | | | |
| KWMTBOMO11304 (BGIBMGA001988) | | | PREDICTED:_transient_receptor_potential_cation_channel_trpm_isoform_X1_[Bombyx_mori] | |
| Gene Information · Genome Browser · Gene Ontology and Pathway · Transcriptional Analysis · View Epigenomics Data · Protein Structure · Population genetics | | | | |
| KWMTBOMO12214 (BGIBMGA014186) | | | PREDICTED:_transient_receptor_potential_channel_pyrexia-like_isoform_X2_[Bombyx_mori] | |
| Gene Information · Genome Browser · Gene Ontology and Pathway · Transcriptional Analysis · View Epigenomics Data · Protein Structure · Population genetics | | | | |

B.

| | |
|---|---|
| Gene | **KWMTBOMO08141** Validated by peptides from experiments |
| Pre Gene Modal | BGIBMGA001085 |
| Description | PREDICTED:_transient_receptor_potential-gamma_protein_[Bombyx_mori] |
| Location | **Bomo_Chr13(-):17500270-17517839** |
| Full name | **Transient receptor potential-gamma protein** |
| Alternative Name | **Transient receptor potential cation channel gamma** |
| Location in the cell | **PlasmaMembrane Reliability : 3.028** |

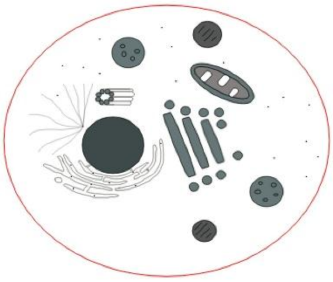

### Sequence

CDS

```
ATGCTGTTGCAACAGCTGAGCAAAAGCGGCGCCTCCGAGATGTCCGGCGGCAGTGGCGGCGGCGGCTCCAGTGCTGACGGGCGGCGTCCCTCCACCTGCCGCGTGGACATGGGCGCGC
TGCTGGCGTCCGAGCCGCGCCCGCTCGACAGCAAGGTTAAGCGGCACAGCATCCACGGGATGACGGAAGAAGAAAATGTGGTTCGACCACATCAGGAGATGGCCGTGCTGTCTTTGGA
GGAGAAGAAGTACCTGCTGGGCGTGGAGCGAGGAGACGTGGCGGGGACGCGGCGCGTGCTGCAGCGGGCCCGCGACACCGGACACATCAACGTCGACTGTGTGGACCCGCTCGGACGC
AGCGCGCTGCTCATGGCCATTGACAACGAGAACCTCGAGATGGTGGAGCTGCTCCTCGAGTTCGGCGTGGAGACCCGCGACGCACTGCTGCACGCCATCTCCGAGGAGTTCGTCGAGG
CCGTGGAGGCCCTGCTCGACCACGAGGAGCGGACCAGGAAGCCCGGCGAACCGAACAGTTGGGAGGCGCTTCCGCCGGAGACGGCCACGTTTACGTCTGACATCACGCCGCTGATCCT
GGCGGCGCACCGGGACAGCTACGAGATCATCAAGCTGCTGCTGGACCGCGGCGCGCAGCTGCCCGAGCCGCACGACGTGCGCTGCGGCTGCGACGACTGCGTGCGCTCCCGCCGCGAG
GACTCGCTGCGACACTCCCGCTCCAGGATCAACGCGTACCGCGCGCTCGCCTCCCCTTCGCTCATCGCCCTATCCTCCAAGGACCCCATTTTAACCGCTTTCGAGCTGTCGTGGGAGC
TGCGCCGGTTGTCCGCATTGGAACATGAGTTCAAAACGGAATACCAGGAGCTCCGCGTTCAGTGTCAGGAGTTCGCGACTGCACTGTTGGATCACACGCGGACGTCTAATGAACTTCA
GATCCTGCTGAATCACGAGAAGGGTGCTTCACCTCAGGCACCGCTCACTGAACCAGGCGCTCCCGAGCGGATGCGCTTGTCGCGGCTCAAGCTTGCCATCAAGTTGCGACAAAAGAAG
TTCGTGGCGCACCCGAACGTGCAGCAGCTGCTGGCGTCGATCTGGTACGAGAGCGTGCCCGGGTTCCGCCGCAAGAACATGCTGCTGCAGGCGGCGGAGATGGTGCGGATCGGCGCCA
TGTTCCCGCTCTACTCGCTCGCCTACATCGCGGCGCCGCACTCCGCCGTCGGACGCACGCTCAGGAAGCCCTTCATCAAATTCCTCTGCCACTCCGCCAGTTACTTTATGTTTCTATT
TCTGTTGATCCTGGCGTCGCAGCGCATCGAGACGGCGCCGGGCGGGCTGCTGTGGGACGTCTCCCACGACGAGCCGCTCTCCCGGCGCGGCTCCATGCCCTCCATAGTCGAGTGGCTC
ATACTCGCCTGGGTCAGCGGCTTAATCTGGAGCGAAGTGAAGCAGCTGTGGGACATGGGGCTGAGAGAGTACGTACATGACATGTGGAACGTCATCGACTTCGTCACCAACTCGCTGT
ACGTGGCCACCGTGGCGCTCAGGATCGTCTCGCACTACCAGGTTCGTCGCGAGATGGCGATGGGCCTGCAGTGGAACCAACCTCGCGAGAAGTGGGACGCTTGGGACCCGATGCTGCT
CTCCGAGGGGCTGTTCTCGGCCGCCAACATCTTCAGCAGTCTCAAGTTGGTGTACATCTTCAGCGTGAACCCTCACCTCGGTCCGTTGCAGGTGTCGCTGAGTCGGATGGTGTTGGAC
ATCCTCAAGTTTTTTGTCTTGGACATACTCGTCATATTTGCGTTCTCATGCGGGCTAAACCAGTTGTTGTGGTACTACGCGGACATGGAGAAGAAGCGCTGCACGGTGGGCACCGCGC
TCACGCCCAACGGCACCCTGCCCGACCCGGACGCCTGTGTCGTCTGGAGACGGTTTGCTAATTTGTTCGAGACAATGCAGACGCTGTTCTGGGCTGCGTTCGGGCTGGTGGACTTGGA
TTCGTTCGAGCTCGACGGGATCAAGATCTTCACGCGGTTCTGGGGGATGCTCATGTTCGGCACGTACGCCGTCATCAACGTCATCGTGCTGCTCAACTTGCTCATCGCCATGATGAAC
CACTCCTATCAGCTGATATCCGAGCGTGCAGATGTGGAATGGAAGTTCGCCCGCAGCAAGCTGTGGATCAGCTACTTTGAGGAGGGTGGAACTGCGCCCCCGCCCTTCAACGTGCTGC
CCTCGCCCAAGTCCCTGCTGTACGCCTGGCGGTGGCTGCAGCGCCGGCTCTGTGGGCACGCCCGCGCCAAGCGAGAACATATGCGGACTATTAGGAGAAAGGTGAAGCAAGCCAATGA
GAGAGACTTTCGATATCAGGCTATAATGCGGAACCTGGTGCGTCGCTACGTGACGGTCCAGCAGCGGCGCGCGGAGAGCGGCGGCGTCACCGAGGACGACGTCAACGAGATCAAGCAG
GACGTCTCCGCCTTCCGCTGCGAGCTCGTCGAGATCCTGCGCAACTCCGGGATGAACACCTCCACAGCCAACGCGGGAGCTCCCGGCGGCGGAGGCGGCAAAAAGAATCGTCAGAAGG
AGCGACGCCTCATGAAGGGATTCAACATAGCTCCGGGCGGGTCCCTGGCCCCCGTGGACGAGTTCATGTCGCCCGTGTCCTGGCTGCAGCACGACCACGGCGTGCCGCACTACTCGCT
GTCCACGCTGCTCGGGCCGCGCCTGCGAGCCTCCCAGTCCAGCCTCAGCGACGGACCCGGCGCCGGGATGAGTGCGTCGCGTCGCAAGCCGCAGCACAAGCGTCGCTGGGGCACAATC
ATCGACGCCGCTCGGGCGGCCCGTGTGTCTCGACTCATCGGCCGGTCGCGCTCTGAGGACTCTGTCTGCGATCACGCGCGGCAGTCGCCAAGCGGTAGCGAGCCTTCGGAGTCGGGCA
GCGACTCGCAGCGCAGTCCGGAGCGGACGGGATCTCGGAGTGGACCCTTGCATCCGTTGACGGCACTGGCGGCCCTCAAGCGGAAGCGAAAGAAATTCTCGGATTCCCGGCGGCCCGA
AGCCACGGTCCGGCCCGTTCCGGCGCCTGAGAGTCTGCAGCGCGCCAGTTCGGTGCCGGCTCGCGTGCCCCCCGCCGCCCCGCCCCGCGCCCCTCCGCCGCCCGCCGTGTCGCCCTCC
ACCACCGCCGAGAGCGTGGCCGCGCGCGGAGGCAGCCGGGAGCCGCTGCTGGCCAGCCTCGACGACGAAGCGCACAAGCGGGACGGCTGCGGCCAGTGCGGGTACGAGTCGGGGCCGC
AGGGCGGCGGGGAGGCGGGGGGCGGGGCGGGCGAGGCGGAGGGCGCGTGCGGGCGGTGCGCGGCGCTGGCGCGGCTGGCGGGCGTGACGCCGCTGGCGGGCCACGCGCCGCACCACTC
CGCGGGCTGGCTCTAG
```

C.

### Summary

| | |
|---|---|
| Function | A light-sensitive calcium channel that is required for inositide-mediated Ca(2+) entry in the retina during phospholipase C (PLC)-mediated phototransduction (By similarity). Forms a regulated cation channel when heteromultimerized with trpl. |
| Subunit | Interacts preferentially with trpl and interacts to a lower extent with trp. |
| Similarity | Belongs to the transient receptor (TC 1.A.4) family.<br>Belongs to the transient receptor (TC 1.A.4) family. STrpC subfamily. |
| Keywords | ANK repeat Calcium Calcium channel Calcium transport Complete proteome Ion channel Ion transport Membrane Reference proteome Repeat Sensory transduction Transmembrane Transmembrane helix Transport Vision |
| Feature | chain Transient receptor potential-gamma protein |
| Uniprot | H9IV05 A0A212EVK1 A0A2H1WAA9 A0A2A4J9Y4 A0A2W1BXK2 A0A182RRP8 + More |
| Pubmed | 19121390 22118469 28756777 12364791 20966253 26227816 + More |
| EMBL | BABH01000036 BABH01000037 BABH01000038 AGBW02012175 OWR45522.1 ODYU01007315 + More |
| Proteomes | UP000005204 UP000007151 UP000218220 UP000075900 UP000075920 UP000075902 + More |
| PRIDE | H9IV05 B7YZW4 Q9VJJ7 |
| Pfam | PF08344 TRP_2 + More |
| Interpro | IPR020683 Ankyrin_rpt-contain_dom + More |
| SUPFAM | SSF48403 SSF48403 + More |
| Gene 3D | 1.25.40.20 1.20.1560.10 |
| CDD | cd00204 ANK |
| oteinModelPortal | H9IV05 A0A212EVK1 A0A2H1WAA9 A0A2A4J9Y4 A0A2W1BXK2 A0A182RRP8 + More |
| PDB | 5Z96 E-value=0, Score=1787 |

### Ontologies

| | |
|---|---|
| KEGG | 101736324 K04967 transient receptor potential cation channel subfamily C member 4 \| (RefSeq) transient recepto |
| PATHWAY | 04745 Phototransduction - fly - Bombyx mori (domestic silkworm) |
| GO | GO:0005262 GO:0016021 GO:0042626 GO:0005524 GO:0006811 GO:0016020 GO:0070588 GO:0051480 GO:0015279 GO:0070679 GO:0005887 GO:0006828 GO:0034703 GO:0005261 GO:0006812 GO:0007628 GO:0043025 GO:0009416 GO:0050908 GO:1990635 GO:0050884 GO:0006816 GO:0008381 GO:0016028 GO:0016740 GO:0005515 GO:0005216 GO:0015914 GO:0006412 GO:0016876 GO:0043039 GO:0055085 |
| PANTHER | PTHR10117 |

D.

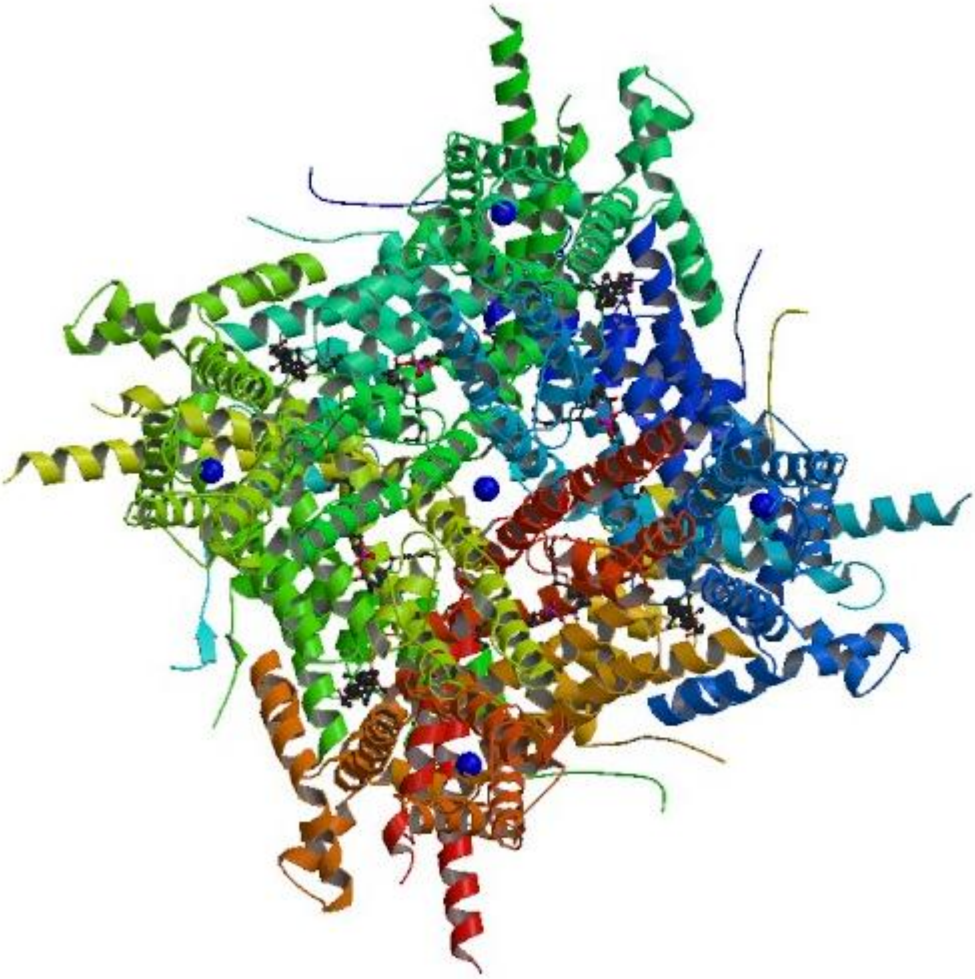

E.

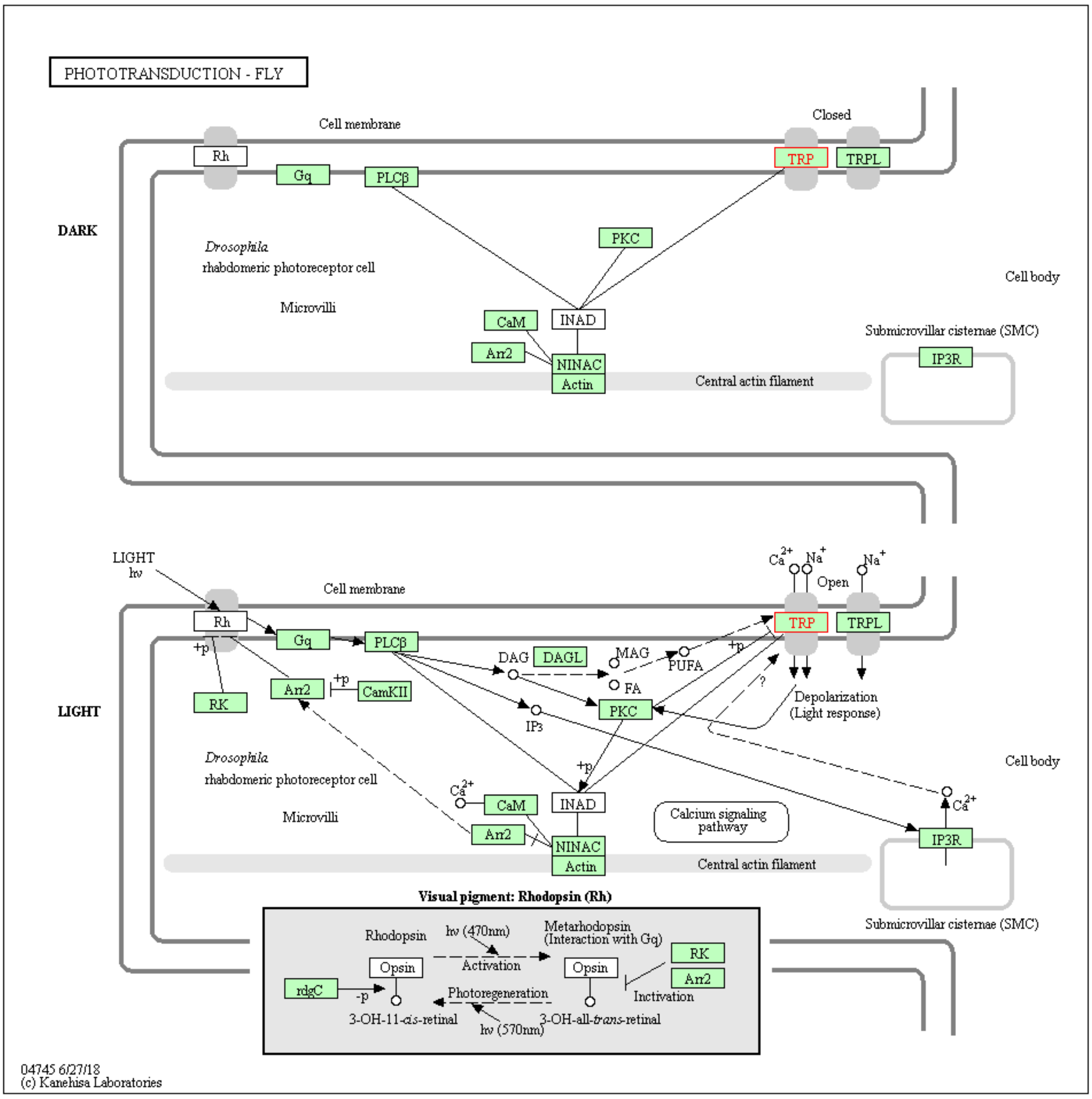

PHOTOTRANSDUCTION - FLY
Cell membrane
Closed
Rh
Gq
PLCβ
TRP
TRPL
DARK
PKC
Drosophila
rhabdomeric photoreceptor cell
Cell body
Microvilli
CaM
INAD
Submicrovillar cisternae (SMC)
Arr2
NINAC
IP3R
Actin
Central actin filament
LIGHT
hν
Open
Cell membrane
+p
DAG
DAGL
MAG
PUFA
FA
RK
Arr2
CamKII
IP3
Depolarization
(Light response)
LIGHT
Calcium signaling pathway
Visual pigment: Rhodopsin (Rh)
Rhodopsin
hν (470nm)
Metarhodopsin
(Interaction with Gq)
Opsin
Activation
Opsin
RK
Arr2
rdgC
-p
Photoregeneration
Inctivation
3-OH-11-cis-retinal
hν (570nm)
3-OH-all-trans-retinal
04745 6/27/18
(c) Kanehisa Laboratories

F.

### Topology

| | |
|---|---|
| Subcellular location | **Membrane** |
| Length: | **1145** |
| Number of predicted TMHs: | **6** |
| Exp number of AAs in TMHs: | **130.89921** |
| Exp number, first 60 AAs: | **0** |
| Total prob of N-in: | **0.00796** |

| | |
|---|---|
| outside | **1 - 388** |
| TMhelix | **389 - 411** |
| inside | **412 - 417** |
| TMhelix | **418 - 440** |
| outside | **441 - 499** |
| TMhelix | **500 - 522** |
| inside | **523 - 549** |
| TMhelix | **550 - 572** |
| outside | **573 - 593** |
| TMhelix | **594 - 616** |
| inside | **617 - 681** |
| TMhelix | **682 - 704** |
| outside | **705 - 1145** |

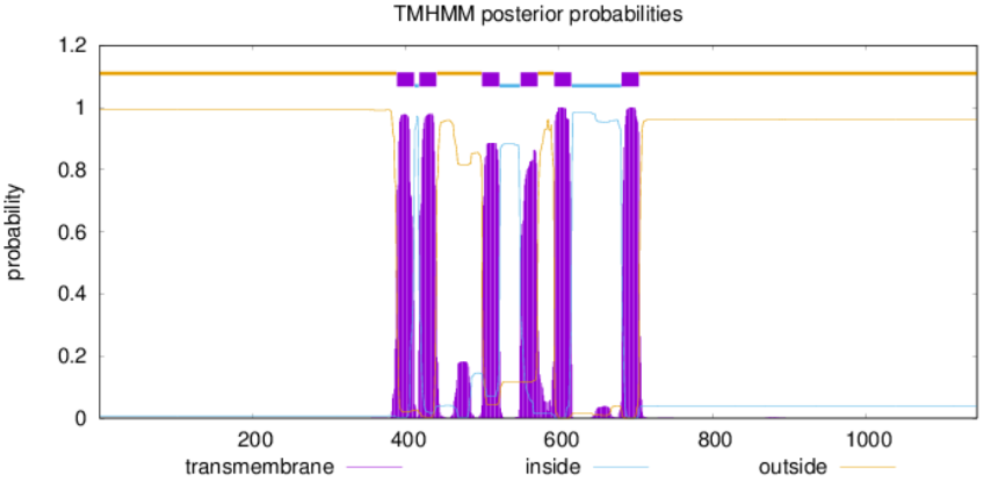

G.

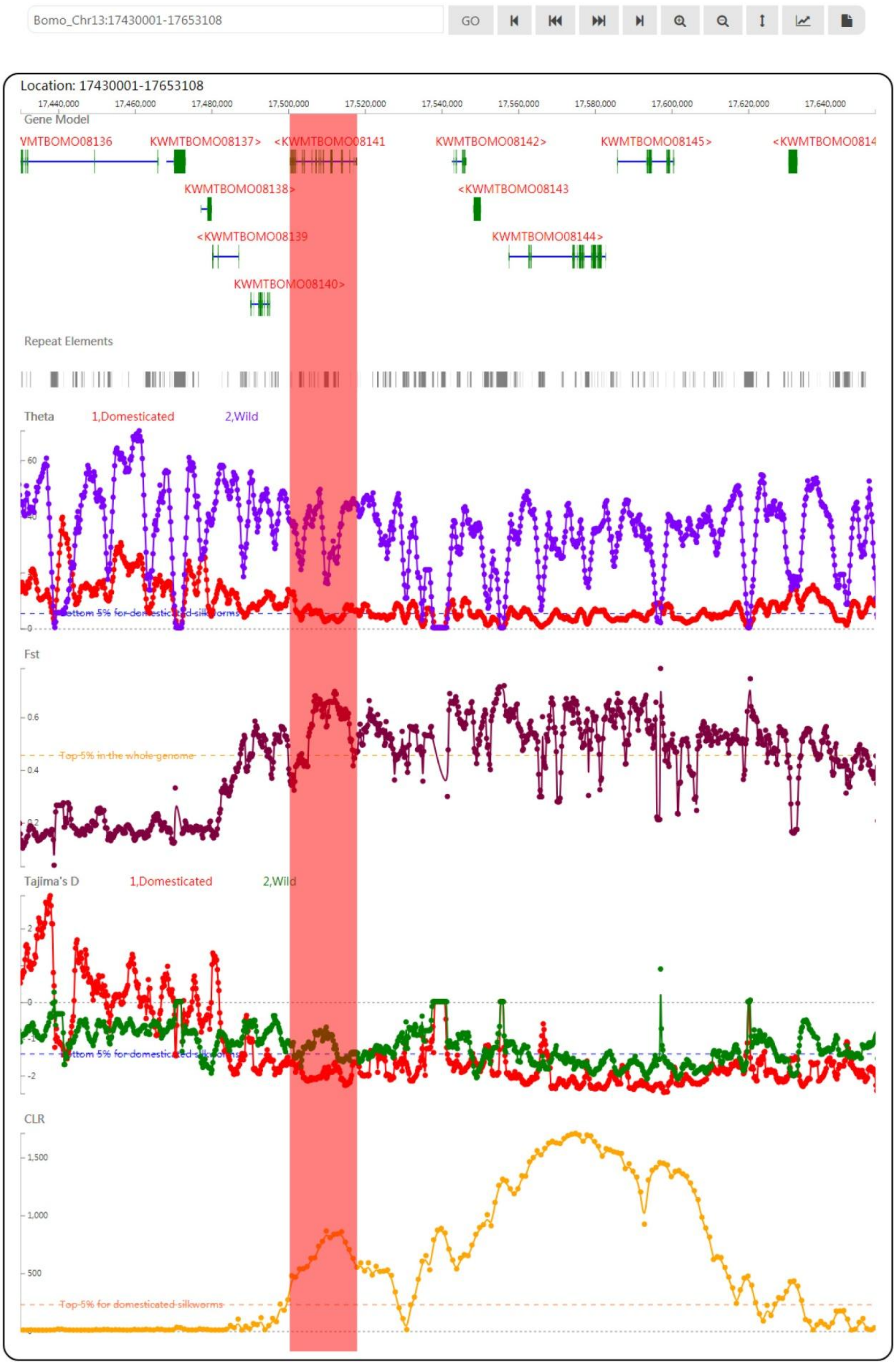

Bomo_Chr13:17430001-17653108
GO
Location: 17430001-17653108
17,440,000
17,460,000
17,480,000
17,500,000
17,520,000
17,540,000
17,560,000
17,580,000
17,600,000
17,620,000
17,640,000
Gene Model
VMTBOMO08136
KWMTBOMO08137>
<KWMTBOMO08141
KWMTBOMO08142>
KWMTBOMO08145>
<KWMTBOMO0814
KWMTBOMO08138>
<KWMTBOMO08143
<KWMTBOMO08139
KWMTBOMO08144>
KWMTBOMO08140>
Repeat Elements
Theta
1,Domesticated
2,Wild
60
40
0
Bottom 5% for domesticated silkworms
Fst
0.6
0.4
0.2
Top 5% in the whole genome
Tajima's D
1,Domesticated
2,Wild
2
0
-1
-2
Bottom 5% for domesticated silkworms
CLR
1,500
1,000
500
Top 5% for domesticated silkworms

H.

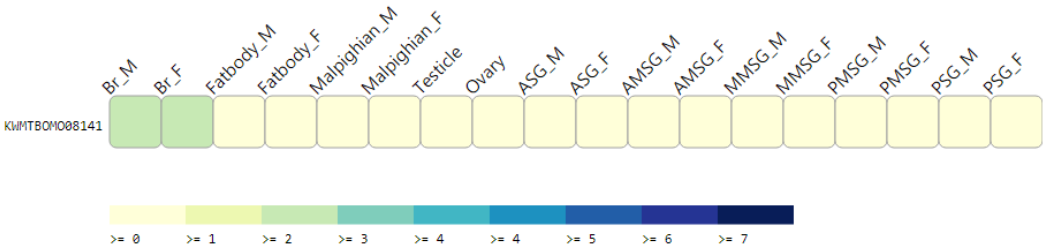
Br_M
Br_F
Fatbody_M
Fatbody_F
Malpighian_M
Malpighian_F
Testicle
Ovary
ASG_M
ASG_F
AMSG_M
AMSG_F
MMSG_M
MMSG_F
PMSG_M
PMSG_F
PSG_M
PSG_F
KWMTBOMO08141
>= 0
>= 1
>= 2
>= 3
>= 4
>= 4
>= 5
>= 6
>= 7

I.

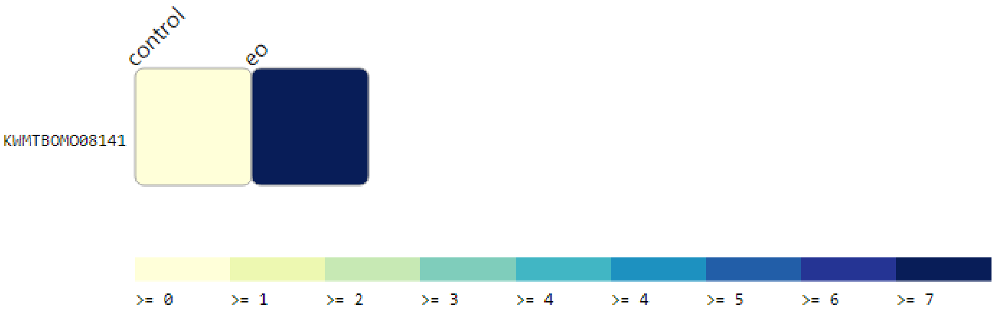
control
eo
KWMTBOMO08141
>= 0
>= 1
>= 2
>= 3
>= 4
>= 4
>= 5
>= 6
>= 7

J.

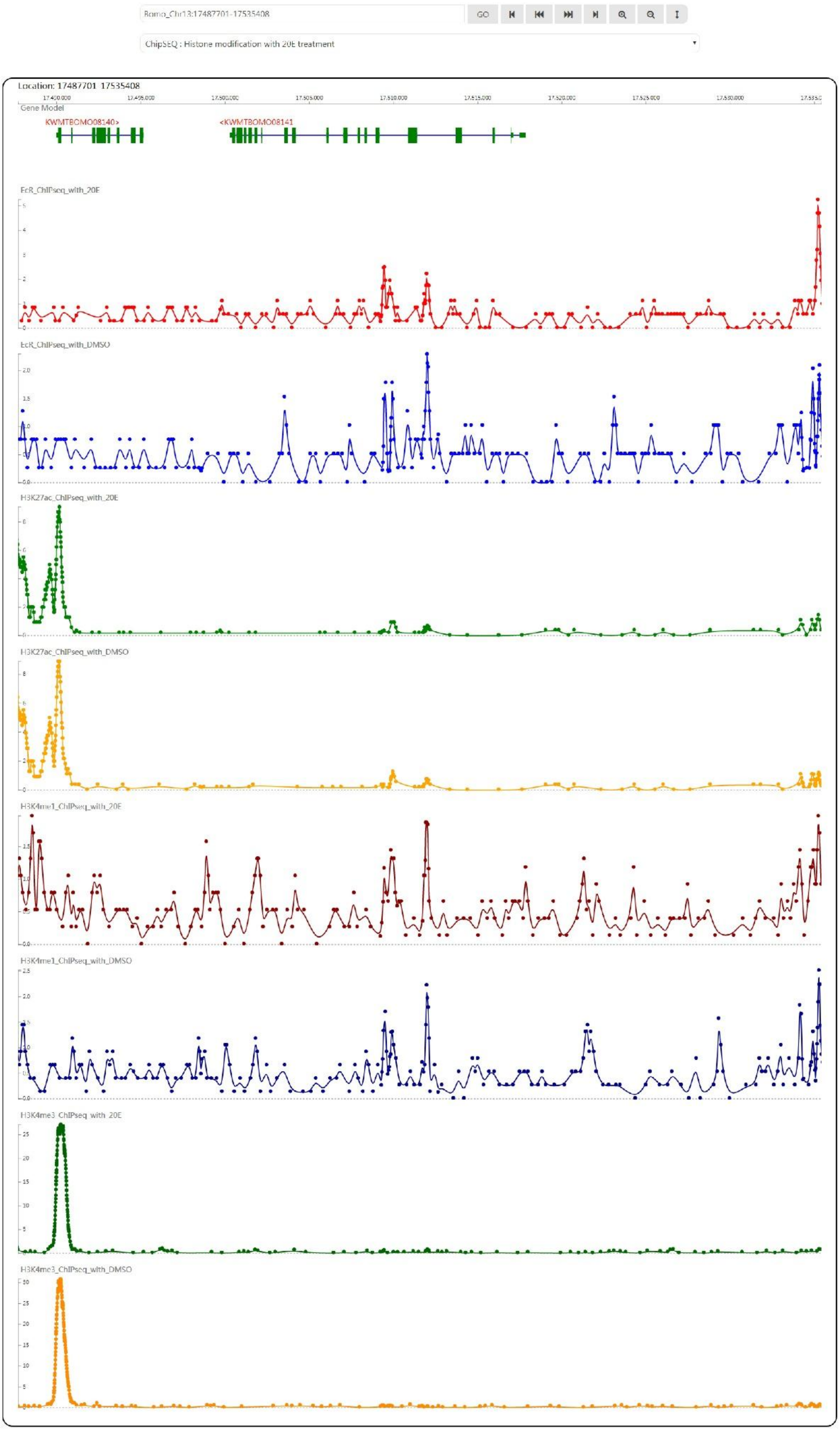

Bomo_Chr13:17487701-17535408
GO
ChipSEQ : Histone modification with 20E treatment
Location: 17487701-17535408
Gene Model
KWMTBOMO08140>
<KWMTBOMO08141
EcR_ChIPseq_with_20E
EcR_ChIPseq_with_DMSO
H3K27ac_ChIPseq_with_20E
H3K27ac_ChIPseq_with_DMSO
H3K4me1_ChIPseq_with_20E
H3K4me1_ChIPseq_with_DMSO
H3K4me3_ChIPseq_with_20E
H3K4me3_ChIPseq_with_DMSO

Figure S1. An example showing the online analysis of KWMTBOMO08141 in SGID. A is a snapshot of the search result for “transient receptor”. The result page lists correlated gene followed by links to gene details, genome browser, gene ontology and pathway, transcriptional analysis, epigenomic signal visualization, protein structure and population genetics analysis results. B is a snapshot of the gene detail page. Subcellular location of the gene is colored in red in a diagram following the basic information. C is the “summary” and “Ontology” parts in the detail page. D is the protein structure of 5Z96, which is of significant similarity to KWMTBOMO08141. E is an online pathway analysis result provided by SGID. F is a snapshot of the topology prediction in SGID. G is a zoom out view of the genome browser with KWMTBOMO08141 marked by a focus bar colored in red. H is the expression of the gene in different tissues (PRJNA284192). M, male. F, female. Br, brain. ASG, anterior silk gland. AMSG, anterior middle silk gland. MMSG, middle middle silk gland. PMSG, post middle silk gland. PSG, post silk gland. I is the expression of the gene in the middle silk gland (PSG) of normal (control) and ecdysone oxidase overexpressed (eo) samples (PRJNA272381). J. A view of epigenomic signals at KWMTBOMO08141 in 4 cell line (EcR, H3K27ac, H3K4me1 and H3K4me3) treated with DMSO or 20E (PRJNA450142).
